## Supplementary material for "Fog signaling has diverse roles in epithelial morphogenesis in insects": Primer list

| | |
|---|---|
| Tc-T48 (1)_F (TC033934) | GCCCCAAAGGATGTATCAAA |
| Tc-T48 (1)_R (TC033934) | GAGTGGCATGAAGTGCAGAA |
| Tc-T48 (2)_F (TC033934) | TTATAACACTGAGGCCCGCA |
| Tc-T48 (2)_R (TC033934) | CTTTGGGGCCACATTTCAGG |
| Tc-mist (1)_F (TC010654) | GCCTTCTTCTGGCTCAACAC |
| Tc-mist (1)_R (TC010654) | AAGACCCTTGGAAGCAGTT |
| Tc-mist (2)_F (TC010654) | ATGACGGCTGTTAGGGTGAA |
| Tc-mist (2)_R (TC010654) | GACACAGCGATTGAGAAGCA |
| Tc-fog (1)_F (TC006723) | TAATCACCGTTGCATCTTGC |
| Tc-fog (1)_R (TC006723) | ATCGTGGTCACTTCCACCTC |
| Tc-fog (2)_F (TC006723) | CCAGTACAGAAGGTGGGGAG |
| Tc-fog (2)_R (TC006723) | ACCTAACAGTCGAACAAAAGGT |
| Tc-cta (1)_F (TC034430) | TCAAGCTGTGGCGGGATAG |
| Tc-cta (1)_R (TC034430) | CAGTGGATGTTCGTCTCGGGATT |
| Tc-cta (2)_F (TC034430) | AATCCCGAGACGAACATCCA |
| Tc-cta (2)_R (TC034430) | TTACATAAGAACCCGGCGGA |
| Of-mist_F | CTTGACAAAAGCCAGGAAGC |
| Of-mist_R | GCGCACATTACCATTGTCAC |
| Of-cta_F | GAACAGGTTGAGTTGTGGGG |
| Of-cta_R | AGTGGCTTATTTGGCTCCCT |
| Of-fog_F | AGTGTTCTCATCCCCGTGTC |
| Of-fog_R | GTCGGCTCAACCGTTGTACT |
| Gb-cta_F | CAG AAG CGG AGC TAG GC |
| Gb-cta_R | GAA TGC TGT CTG GCA CAT AAC |
| Gb-mist_F | CAT CTG GTG GAC CTt CAG AG |
| Gb-mist_R | CAT TCA TTA GCA TGC GTA AGT G |
| Gb-fog_F | CTGTCGACGCCAGACGCTC |
| Gb-fog_R | GTGGAGGAGGTGGGCAGT G |
| Tc-tap_F (TC034722) | CGGGAATACGCCTAAGTTGG |
| Tc-tap_R (TC034722) | TCTGAGAGGCATGTCGTTCG |
